# IDR searcher: a search engine solution for public image resource

**DOI:** 10.64898/2025.12.16.694566

**Authors:** Khaled Mohamed, William Moore, Dominik Lindner, Josh Moore, Jason R. Swedlow, Petr Walczysko, Frances Wong, Jean-Marie Burel

## Abstract

We introduce the IDR searcher, an open-source search engine designed to ease the exploration of datasets hosted in public bioimaging resources. The application offers a fast, efficient, cost-effective solution for datasets discovery and has the potential to address current disparities in finding quality datasets for exploratory research and to provide the foundation for cross-resource search.

## Introduction

Bioimaging is a dynamic field producing quality imaging datasets, many of which have been made publicly accessible as part of a strong drive to make the data Findable, Accessible, Interoperable and Reusable (FAIR) (Wilkinson *et al*, 2016). Public image data resources are now available, but finding quality data quickly and efficiently remains a challenge, and this in turn limits their reuse and reduces the chance of new discoveries.

In 2016, the Open Microscopy Environment (OME) began a collaboration with EMBL-EBI to build the Image Data Resource (IDR) (Williams. et al, 2017), an added-value, journal-independent resource, publishing reference bioimage datasets associated with peer-reviewed publications.

IDR uses, as its basis, Bio-Formats (Linkert *et al*, 2010) and OMERO (Allan *et al*, 2012).

Bio-Formats is a suite of libraries which are used heavily by the Java image processing community for reading proprietary scientific image data and metadata into a common model. OMERO is a secure client-server software platform for image data management and analysis. IDR uses the OMERO database to store image metadata and a collection of web plugins based

### on OMERO.web to explore and display the data and metadata associated with datasets it publishes

Since the creation of IDR, other public bioimage data resources have emerged to collect and share data, including analytical results such as the BioImage Archive (BIA) (Hartley et al, 2022) and SSBD (Tohsato et al, 2016).

These resources are now hosting over one PetaByte of well-annotated, organised, public bioimage data. When initially built, the emphasis was put on data access either via User Interface (UI) or Application Programming Interface (API), to partially address the FAIR principles, specifically the principles of Accessible (A) and Interoperable (I) data.

For each study in IDR, image data is stored along with structured and unstructured metadata related to the experimental design, data acquisition and analysis. OMERO’s database has specific tables for structured acquisition metadata whereas flexible key-value annotations are used to store the less structured experimental metadata

Currently there is some diversity in the metadata standards and ontologies used by various public bioimage data resources.. Not all metadata (e.g. structured metadata) is readily available to standard searchers (Google, Bing, etc.) for indexing. The combination of these challenges mean that datasets that reveal key insights into biological mechanisms and diseases cannot be easily found, thus limiting sharing, impact and reuse.

Over two decades ago, open-source libraries offering full-text search and indexing of data emerged, with Apache Lucene (https://lucene.apache.org/) being one of the flagship tools. These libraries quickly became the backbone of several enterprise search servers, which index either structured or unstructured data from different sources associated with a specific repository, e.g. a company’s data or data from a specific scientific domain. Two heavily adopted open-source options, both built on Apache Lucene, are Apache Solr (https://solr.apache.org/), and Elasticsearch (https://www.elastic.co/). Apache Solr is the solution behind the search capabilities of the Protein Data Bank (PDB) (Berman et al, 2000), and Elasticsearch is used by the Netflix recommendation engine^1^ and the eBay large scale search functionality^2^.

In this paper, we introduce a resource-agnostic searching solution leveraging Elasticsearch to make bioimaging data hosted across public resources easily and quickly findable (“F” in FAIR)

We describe the use of ElasticSearch to meet the indexing and search requirements of metadata-rich bioimaging repositories like IDR.

## Results

We developed the IDR searcher (https://github.com/ome/omero_search_engine), an open-source search engine that leverages the full-text search and specifically the lexical search capability of Elasticsearch. The IDR searcher offers a fast, efficient, cost-effective solution to explore large volumes of data stored in different resources; thus, it provides a viable, performant, scalable search solution for the bioimaging community.

### Architecture

The IDR searcher is composed of three components described in the schematic architecture diagram (Fig. 1).

- **Data indexing**: We have designed the search engine so that it is independent of the underlying source of data. The search engine can index data stored in diverse structures, such as relational databases, database dumps or text-based files.
- **API access**: The search engine prioritises the developers’ experience with a JSON-centric REST API. JSON is the standard format for transferring data.
- **Horizontal scaling**: The search engine is designed to work with several Elasticsearch nodes to guarantee high availability and performance.

**Figure 1.**
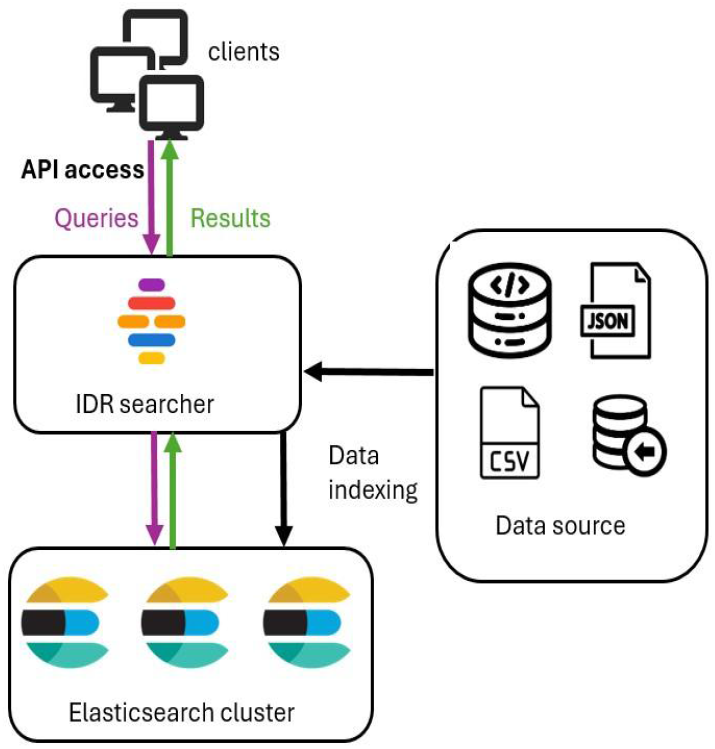
This diagram represents a schematic view of the architecture

### Data Indexing

When using the IDR searcher, one first has to index the data and build a cache which provides an overview of the indexed data, for example, the number of images linked to a specific organism. Data is first retrieved from a source such as a relational database. The data is then formatted according to the indexer template (see https://www.elastic.co/docs/manage-data/data-store/templates) before being pushed to Elasticsearch via its API. The cache is then built when the indexing is complete.

The architecture of the search engine is implemented to be data source agnostic and is therefore able to be used to index and search publicly available metadata stored in other resources. We encapsulate information about the original data source in the indexed data (data provenance). This key point will enable the integration, within a single search index, of data originating from diverse resources like the IDR or the BioImage Archive; an idea that we are currently exploring. This represents a step towards a federated search solution enabling users to query independent bioimaging resources through a common search interface.

Data is being continuously added to bioimaging resources. In the IDR, this was leading to increased indexing and caching times on each release date. To address this issue, we prioritised our effort on reducing the indexing and caching workload to avoid delays in publishing new datasets. (see Fig. 2).

**Figure 2.**
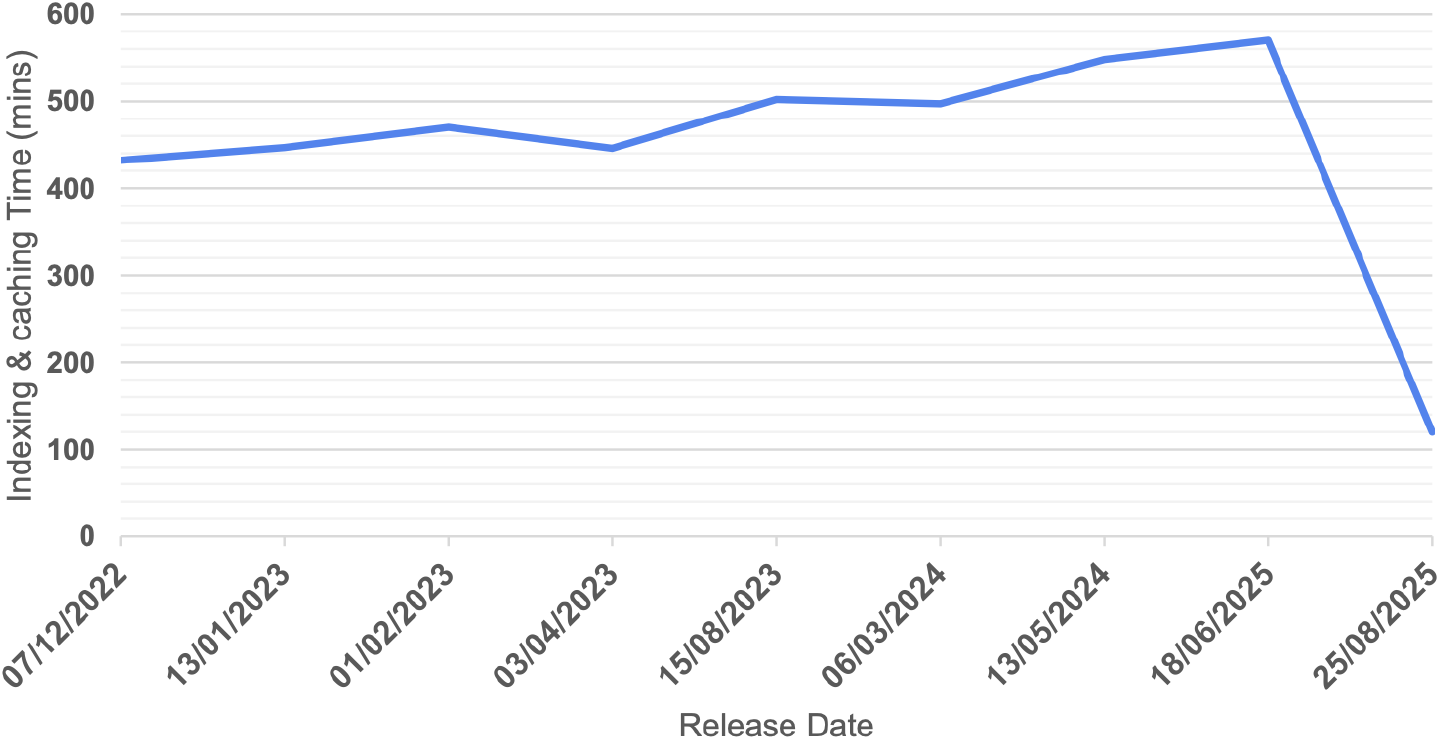
The graph shows the time needed for the indexing and caching of publication releases within IDR. The total image count was 13.1 million on December 7th, 2022, increasing to 14 million by August 25th, 2025. During the August 25th, 2025, release, only the newly added study was indexed and cached, contrasting with previous releases where all images hosted in IDR were indexed and cached.

We enhanced the search engine to only index newly added or updated data. This improvement, implemented with the publication on August 25th, 2025 of a new study composed of 221 images, reduced the pre-publication indexing and caching time by approximately 7 hours. By refining our current caching strategy, we anticipate further time reductions.

### API access: searching the data

The IDR searcher exposes a JSON-centric REST API over HTTP and provides a collection of documented endpoints (https://idr.openmicroscopy.org/searchengine/apidocs/), allowing front-end applications to easily access the indexed data. It acts like a broker that internally uses the Elasticsearch API to construct advanced queries composed of three steps, defining the search context:

1. **Search scope:** This establishes the search’s scope, for example, whether to search for images across the entire repository or within a specific study.
2. **Key-Value Search:** This defines an optional key and value to look for within the scope of the search, e.g. Gene Symbol contains PAX 6.
3. **Search condition:** This option allows the addition of further search conditions, e.g. Gene Symbol contains Pax 6 AND Imaging Method equals SPIM.

Upon receiving a query from a client application, the search engine queries Elasticsearch via its API, then undertakes the task of converting the raw output from Elasticsearch into a structured JSON format. This JSON output is then returned to the client application (see Fig. 1).

The returned JSON structure provides a comprehensive overview of the search operation; it encapsulates key information, including the original data source, the actual search results, a clear representation of the query that was executed, the total number of results found, and the time taken to perform the query. Such information helps to correctly handle the results in client applications. The search engine also supports pagination (results are divided into multiple numbered pages), which is critical for efficiently handling large volumes of metadata, preventing potential system slowdowns or failures, and considerably improving the user experience.

We have prepared a collection of notebooks (https://workflowhub.eu/collections/36) hosted in the WorkflowHub (Gustafsson, O.J.R et al, 2025), demonstrating how to use the search engine REST API to explore the richness of IDR.

### Horizontal scaling

We designed the IDR searcher so it can easily scale horizontally using the distributed architecture of Elasticsearch. The search engine works with a cluster of Elasticsearch nodes to guarantee high availability and performance. High availability is achieved by increasing the number of Elasticsearch nodes and high performance is achieved by deploying nodes on separate machines. Increasing the number of nodes requires minimal configuration.

### Deployment

The search engine can be deployed using either Docker, a software platform to build and deploy applications or Ansible, an automation tool to configure, manage and deploy applications. The Ansible role is open-source and publicly available at https://github.com/ome/ansible-role-omero-searchengine. The instance of the search engine deployed alongside IDR is managed by Ansible playbooks on an OpenStack-based cloud contained within the EMBL-EBI Embassy resource. It is configured to use three Elasticsearch nodes deployed on the same Virtual Machine. As the volume of data hosted in IDR increases, we plan to deploy each Elasticsearch node to a separate Virtual Machine.

### Use case: Integration with the IDR User Interface

When IDR was first created, we initially developed and used OMERO.mapr (https://github.com/ome/omero-mapr), as the search solution for IDR. We further enhanced the search functionality of OMERO.mapr with a front-end application, IDR.gallery (https://github.com/IDR/idr-gallery), to provide a search User Interface and display search results retrieved from the relational database.

We first deployed both OMERO.mapr and IDR searcher side-by-side so we could compare performance and ensure that equivalent queries using either tool returned identical results. We used a single Elasticsearch node to ensure a fair performance comparison between applications. We performed several queries searching for a specific Gene Symbol, Compound Name or Phenotype.

Fig. 3 highlights a performance improvement for each query. OMERO.mapr query times are highly variable depending on the query complexity, but in general are between a couple of seconds to a couple of minutes slower than the equivalent query in IDR.searcher. The performance of IDR.searcher can easily be improved by adding more Elasticsearch nodes to the search cluster.

**Figure 3.**
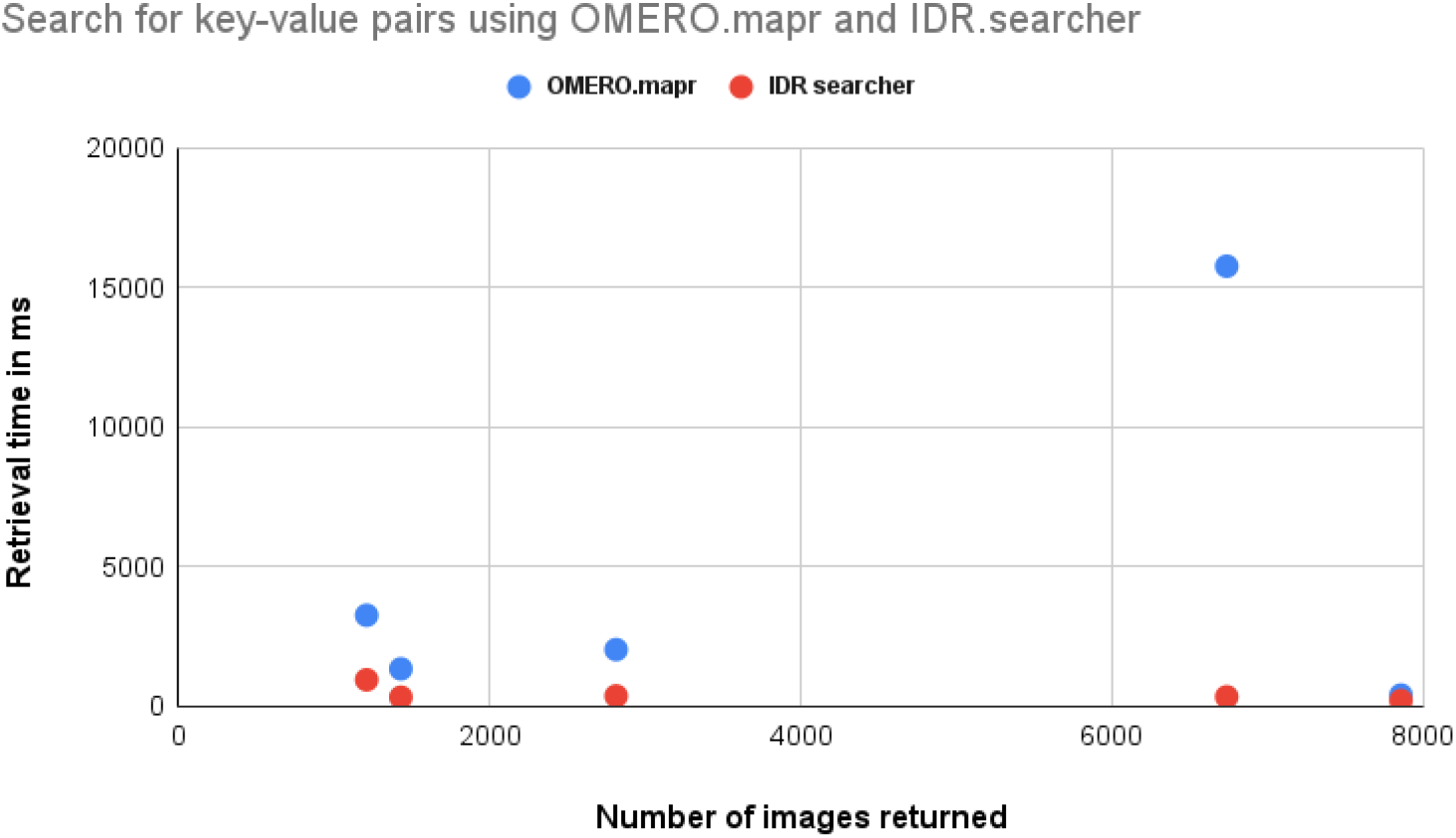
Scatter plot comparing the time to perform a specific query using OMERO.mapr and IDR searcher. For each query, the same images are returned using either application.

### Use case: Integration with BioFile Finder

Studies in added-value resources like IDR usually have large amounts of associated metadata requiring applications whose primary focus is metadata exploration. BioFile Finder (https://bff.allencell.org/) is a Web application from the Allen Institute for Cell Science designed for easy access and sharing of images through metadata search, filtering and sorting. We added to the IDR searcher the ability to export the indexed metadata linked to a study to Comma Separated Values (CSV) files or Parquet Files. Those files can be directly opened in BioFile Finder to offer the user a new way to visually explore, sort, and filter the data and metadata stored in the IDR (see Fig. 4). We have tested the workflows with studies of files composed of over 2 million entries. From our experience, the Parquet format is the most efficient solution for the transfer of large amounts of data to BioFile Finder.

**Figure 4.**
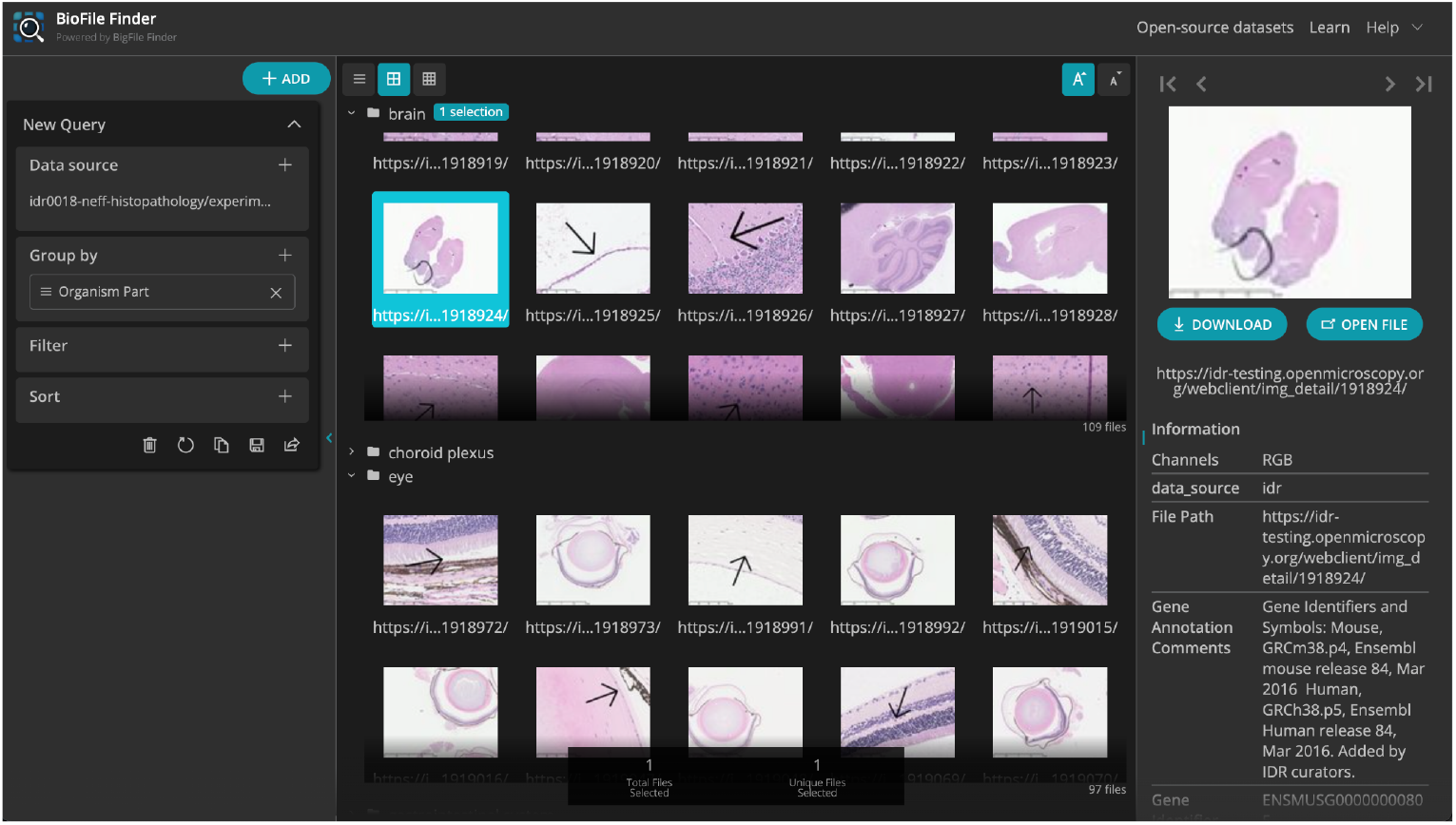
Thumbnails and metadata retrieved from IDR organised in groups in BioFile Finder, grouping data is not supported by IDR.gallery.

This example illustrates the value of bioimage data sharing and shows how the IDR searcher has the potential to contribute to a broader shift toward a more open research culture in bioimaging.

## Discussion

We have developed the IDR searcher to provide an efficient solution for searching through a large volume of public bioimage metadata, which is growing exponentially. The application not only helps finding public bioimage data efficiently but also allows the exchange of metadata between public resources like IDR and metadata visualisation tools like BioFile Finder.

We also aim to take advantage of the IDR searcher resource-agnostic architecture to provide the foundation for searching through the three public bioimage resources, IDR, BIA and SSBD via a single index. This could be achievable by interfacing with the interoperable resource as an outcome of the foundingGide project (https://founding-gide.eurobioimaging.eu/), which is working on harmonising metadata models and mapping ontologies between IDR, BIA and SSBD.

An exciting direction of work will be to provide a common exchange format based on RO-crate (Soiland-Reyes et al, 2022) for exporting indexed metadata associated with studies and outputs of a search. This will not only facilitate the integration with 3rd party tools but could also allow the search outputs to be integrated into OME-Zarr (Moore *et al*, 2021), a flexible, multi-dimensional image format that provides remote access to large multi-TeraByte datasets using Web standards. OME-Zarr has greatly simplified the creation of public resources e.g. https://cryoetdataportal.czscience.com/browse-data/datasets. IDR searcher could also become a major component for searching across those emerging public resources publishing OME-Zarr images.

All those aspects combined will contribute to mitigating the lack of findability and exploration of quality bioimage data.

## Acknowledgements

The authors thank Nathalie Gaudreault, Graham Johnson, Sean Meharry, Daniel Toloudis for valuable comments and discussions on integrating with BioFile Finder. We also thank the foundingGIDE (https://founding-gide.eurobioimaging.eu/) project’s members for the valuable discussions about the architecture of the application. This work was funded by awards from the Wellcome Trust [grant numbers 221361/Z/20/Z, 313803/Z/24/Z] and the European Union’s Horizon Europe research and innovation programme [grant number 101130216]. Views and opinions expressed are, however, those of the author(s) only and do not necessarily reflect those of the European Union or the European Research Council Executive Agency. Neither the European Union nor the granting authority can be held responsible for them.

## Footnotes

1 https://netflixtechblog.com/reverse-searching-netflixs-federated-graph-222ac5d23576

2 https://www.elastic.co/elasticon/tour/2015/’amsterdam/ebay-classified-group-and-a-scalable-flexible-search-platform-to-build-on

